# Screening of herbal molecules for the management of Alzheimer’s Disorder through molecular docking and *in-vitro* investigation

**DOI:** 10.1101/2023.01.01.522412

**Authors:** Priyanka Nagu, Amjad Khan A Pathan, Vineet Mehta

**Author notes:** **Corresponding Author**, Dr. Vineet Mehta, Department of Pharmacology, Government College of Pharmacy, Rohru, Himachal Pradesh, India. **171207**.

## Abstract

At present, there is not a single disease-modifying drug available for the management of Alzheimer’s disease (AD) pathogenesis. The exact pathology of AD is still not known, which opens up the wide scope of research for developing some novel therapeutic strategies for AD management. In the present study, 100 herbal molecules were identified through the literature survey which could be beneficial for Acetylcholinesterase (AChE), Butyrylcholinesterase (BChE), β-Secretase inhibition, and neurodegeneration. AutoDock Tools-1.5.6 docking software is used to screen the herbal molecules against AChE, BChE, and β-Secretase with Protein Data Bank (PDB) ID 1B41, 1P0I, and 1FKN respectively. Based on the docking parameters quercetin, rutin, vitisinol-C, dihydrotanshinone-I, and β-carotene were found to be the best molecules against their respective proteins receptors. Moreover, *in-vitro* AChE and BChE assay demonstrated that quercetin and rutin could modulate cholinergic pathways during AD and thereby could impart beneficial effects during AD. Further, our results for *in-vitro* neurodegeneration studies also support the neuroprotective effect of quercetin and rutin against HgCl_2_-induced neurodegeneration and suggested the protective role of these molecules against neurodegeneration during AD. However, a preclinical investigation is required to support the potential effect on AD pathogenesis.

## 1 Introduction

Alzheimer’s Disorder (AD) is one of the major health care concerns that is characterized by dementia and marked cognitive decline.^1^ Recent time has witnessed a significant increase in the population affected by this lifestyle disorder, especially in developed and developing nations.^2^ World Health Organization has raised serious concerns regarding the increasing global population of AD. As per the World Alzheimer Report 2021, AD and dementia were reported to be the 7^th^ leading cause of global. It was estimated that 57 million people suffered from AD in 2019, besides, it is estimated that the global burden of AD and dementia will increase to 153 million by the year 2050.^3^ AD results from progressive neurodegeneration and thereby loss of normal physiological functioning of the brain, especially in the region of the hippocampus and cortex that are associated with learning, memory, and cognitive functions.^4^ Numerous identified pathways are believed to be directly involved in the development and progression of AD, however, none of them is capable of fully explaining the underlying pathogenesis of AD. Some of these prominent pathways include neurodegeneration, impaired Acetylcholine, cholinesterase functioning, oxidative stress, neuroinflammation, etc.^2^These pathological pathways work in harmony to collectively form a complex interconnected network that contributes to the development and progression of AD.^5^ These pathways are the targets of the currently available AD therapeutic strategy. The majority of the currently available drugs target a single pathway of AD pathogenesis and are aimed at slowing down the progression rate and providing symptomatic relief from the AD with negligible effect on the root cause of AD due to which AD continues to progress throughout life.^6^ Considering the current global burden of Ad and future estimates along with the lack of an efficient therapeutic strategy to completely halt the development and progression of AD, we need to identify better therapeutic interventions that can utilize multiple pathways leading to the development and progression of AD at once to get better management of AD.

Natural sources offer us an enormous amount of therapeutic possibilities for a variety of disorders like cancer, diabetes, CNS ailments, etc. Rivastigmine, atropine, physostigmine, nicotine, muscarine, *Bacopa monari*, are examples where herbal interventions have found clinical applications for the management of CNS complications such as AD, etc.^7, 8^ As evident from the established reports, AD is a result of a complex interplay of different pathological pathways working in harmony, therefore, the best approach to find a better therapeutic strategy against AD would be to identify molecules that could target multiple pathways. The present study was aimed at identifying such molecules that could target multiple pathways leading to AD by utilizing *in-silico* and *in-vitro* techniques.

## 2 Materials and methods

### 2.1 Preparation of the ligands and receptors

In the present molecular docking study, 100 herbal molecules were selected from the literature survey considering their potential to modulate various pathways associated with AD such as acetylcholinesterase (AChE), Butyrylcholinesterase (BChE) pathway, Tau protein phosphorylation, oxidative stress pathway, and inflammatory stress pathway, etc. The 2D and 3D structures of selected herbal molecules were generated through Chemdraw Ultra 7.0 software and Marvin sketch software. We used Auto Docktools-1.5.6 docking software to screen all the herbal molecules against AChE, BChE, and β-Secretase with Protein Data Bank (PDB) ID 1B41, 1P0I, and 1FKN respectively. Selected molecules were converted to pdbqt format by adding gasteiger charges to the ligands. The Auto Dock Tools-1.5.6 docking software is a graphical user interface used for the preparation and analysis of docking results. Receptor preparation was performed according to the method described previously by Trott and Olson (2010)^9^ in which atomic charges, fixed bond, Kolman charges, and hydrogens were added to the proteins and converted to a pdbqt format, which stores the partial charges for further docking studies.

### 2.2 Molecular Docking

The docking was performed with Auto Dock Tools-1.5.6 docking software in which three prepared protein targets, 1B41, 1P0I, and 1FKN, were docked with 100 herbal molecules along with an internal standard. Docking grids were created by adjusting the size of the grid box, which will pick the coordinates within which the docking of ligand and protein was performed. The auto grid and auto dock widgets were used to prepare the docking and grid files.^10^ Finally, docking parameter files and docking log files were generated by running the auto grid and auto dock widgets to check the molecular interaction between ligand and protein molecules. The results of the docking study were estimated in terms of binding energy (kcal/mol) of the different conformations of ligands with the receptors, amino acid interactions, and bond length.^10^To visualize results, PyMOL molecular graphics system and Protein-Ligand Interaction Profiler (PLIP) open-source web server were used.

### 2.3 *In-vitro* evaluation

We screened and identified quercetin, rutin, β-carotene, sumaflavone, and vitisinol-C to be most efficient in targeting 1B41, 1P0I, and 1FKN through docking studies. To narrow down the screening process, these molecules were then subjected to *in-vitro* testing on prominent pathways associated with the pathogenesis of AD. These pathways include neurodegeneration and cholinesterase activity in terms of AChE and BChE activity.

#### 2.3.1 In-vitro Neurotoxicity Assay

The effect of the herbal molecules on the HgCl_2_ induced neurotoxicity was evaluated on Neuro-2a cell lines in a 96-well plate by using 3-(4, 5-dimethyl-thiazol-2-yl)-2, 5-diphenyltetrazolium bromide (MTT) as per the method described previously.^11^ Briefly, cell plates were incubated till 70% confluency in EMEM media. Growth media was replaced with serum-free EMEM media and 10 μl of different concentrations (300 μM) of the herbal molecules was added to each well. An equal volume of phosphate buffer saline (PBS; pH 7.4) was used for the control reaction well. Culture plates were incubated inside humidified CO_2_ incubator at 37°C for 6h. Incubated cells were then treated with 25 μM HgCl_2_ to induce neurotoxicity. Cultures were allowed to grow for 24h inside a CO_2_ incubator at 37°C. Afterward, culture media was removed and cells were washed with PBS thrice. The culture was then incubated at 37°C for 3 h with 100 μl serum-free EMEM having 5 mg/mL MTT. This was followed by cell lysis with DMSO and the absorption of the formazan was recorded spectrophotometrically at 570 nm. Simultaneously, the readings of the control (without treatment) and blank (without any cells). The entire experiment was performed in triplicate and percent neuroprotection was calculated.

#### 2.3.2 Acetylcholinesterase (AChE) and Butyrylcholinesterase (BChE) inhibitory assay

Herbal molecules were further screened for their potential against AChE and BChE enzymatic activity as per the methods described previously^11^, with slight modification. In both of these experiments, donepezil was used as a positive control. Briefly, the stock solutions of the herbal molecules (0.1M) and donepezil were prepared in PBS and stored at 4°C until used. The working dilutions were prepared from the stock solution by proper dilution with PBS. The inhibition of AChE and BChE activity was evaluated in a 96-well plate using 5, 5’-Dithiobis-(2-Nitrobenzoic acid (DTNB) or Ellman’s reagent. Each reaction mixture was composed of 25 μl enzyme solution (AChE or BChE) prepared as 25 mU enzyme in PBS, 75 μl DTNB prepared in PBS (having 6 mM NaHCO_3_), 50 μl PBS, and 25 μl herbal molecule or donepezil (standard reaction) or PBS (control reaction). The reaction mixture was incubated at 37°C for 10 min after which 25 μl of 0.075 M substrate (acetylthiocholine iodide for AChE and butyrylthiocholine iodide for BChE) was added to initiate the reaction. Enzymatic activity was recorded spectrophotometrically at 412 nm over 4 min. Percent inhibition of AChE or BChE activity was calculated using the following equation:

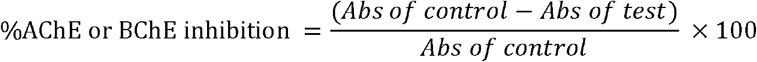

### 2.4 Statistical Analysis

Statistical significance was determined by using GraphPad Prism 6 software. Data were expressed as mean ± SD and the statistical significance was assessed by one-way ANOVA followed by Dunnett’s multiple comparison post hoc test (* p < 0.05, ** p < 0.01, and *** p < 0.001). Duncan’s multiple range test was also performed at p < 0.05.

## 3 Results and discussion

### 3.1 Molecular docking study

The molecular docking technique is one of the most commonly used tools to screen herbal molecules for predicting their interactions with various protein targets. Docking screening provides us with crucial information regarding the interaction of the herbal molecule with the target protein in terms of ligand-protein interaction energy (kcal/mol), bond length, the number of interactions, amino acids involved in the interaction, root mean square deviation (RMSD) values, estimated inhibition constant (Ki), etc., based on which protein specific activity of a test ligand compound was predicted.^10^ Previously, we screened 100 herbal molecules against various targets of AD and predicted the 10 best molecules based on the outcome. The molecules include β-carotene, dihydrotanshinone I, glabridin, liriodenine, morin, N-formylanonaine, quercetin, quercitrin, rutin, and vitisinol C. To get more prominent and confirmatory data for these molecules, in the present study, these herbal molecules were subjected to molecular docking screening against AChE, BChE, and β-secretase (PDB ID: 1B41, 1P0I, and 1FKN respectively) by using AutoDock tools. In the present study, we used Donepezil as an internal standard for all receptors to avoid any variations in the standard for comparison.

During AD, the levels of ACh in the brain are lowered and clinical management of AD includes enhancing ACh levels by inhibiting AChE which results in increasing the ACh levels in the synapse, which provides symptomatic relief from AD and slows down the disease progression.^6^ Inhibiting AChE in the brain is one of the most common therapeutic strategies employed in clinical settings to enhance cholinergic signaling during AD.^6^ The docking interaction results of 1B41 (AChE) are represented in Table 1. In our study, β-Carotene and Viticinol C demonstrated the least binding energy with the target protein with docking scores of −11.45 kcal/mol and −9.63 kcal/mol respectively. These molecules interacted with various amino acids like ARG 292, SER 199, VAL 278, and GLY 117, suggesting their potential to modulate the activity of AChE. The docking score of the standard drug, donepezil, was observed to be −9.44 kcal/mol, which was comparable to dihydrotanshinone I (−9.13 kcal/mol) and glabridin (−9.40 kcal/mol). Interestingly, the binding energy of quercetin with the target receptor was observed to be on the slightly lower side (−7.98 kcal/mol), but it demonstrated interaction with PRO 286, ARG 243, and THR 234, suggesting its strong potential to modulate receptor activity. RMSD values of the herbal molecules were observed in the range of 0.00 to 1.0 and were comparable to donepezil, suggesting good interaction. Moreover, Ki values of the herbal molecules were observed in the range of 4.07 nm to 11.39 mM, and β-Carotene (4.07 nM) and Vitisinol C (87.14 nM) demonstrated better Ki values than Donepezil (202.11 nM).

**Table 1:** Binding energies (kcal/mol), RMSD values, and Inhibition constant (Ki) of selected ligands with PDB: 1B41 (Acetylcholinesterase)

| Ligands | Docking Score (kcal/mol) | Reference RMSD | Estimated Inhibition Constant (Ki) |
| --- | --- | --- | --- |
| $\beta$ Carotene | -11.45 | 0.35 | 4.07 nM |
| Dihydrotanshinone I | -9.13 | 0.00 | 202.11 nM |
| Donepezil | -9.44 | 0.64 | 119.94 nM |
| Glabridin | -9.40 | 0.35 | 128.29 nM |
| Liriodenine | -8.51 | 0.41 | 577.67 nM |
| Morin | -7.65 | 0.64 | 2.48 μM |
| N-Formylanonaine | -2.65 | 0.68 | 11.39 mM |
| Quercetin | -7.98 | 0.74 | 1.42 μM |
| Quercitrin | -7.80 | 0.80 | 1.93 μM |
| Rutin | -7.97 | 1.00 | 1.43 μM |
| Vitisinol C | -9.63 | 0.62 | 87.14 nM |

Along with AChE, recent findings suggest that BChE also plays a crucial role in regulating ACh levels in the brain.^12^ Along with this, BChE is associated with other AD pathways such as β-amyloid plaque deposition, etc.^13^ The docking interaction results of 1P0I (BChE) are represented in Table 2. In this study, the best results were observed for rutin and viticinol C, which demonstrated a docking score of −11.36 kcal/mol and −10.59 kcal/mol respectively. These results were better than the standard that demonstrated a docking score of −10.20 kcal/mol. Moreover, rutin and viticinol C demonstrated strong interaction with amino acids like SER195, GLY 114, THR 117, TYR 125, GLU 194, TYR 329, and GLY 436. Docking interaction of β Carotene, dihydrotanshinone I, glabridin, and quercitrin was observed in the range of −8.89 kcal/mol to −9.66 kcal/mol, which was comparable to the standard drug. RMSD values of the herbal molecules were observed in the range of 0.00 to 0.94, suggesting good interaction. Ki values of the herbal molecules were observed in the range of 4.73 nM to 7.79 mM. Rutin (4.73 nM) and vitisinol C (4.73 nM) demonstrated better Ki values than donepezil (33.18 nM), suggesting that these molecules can efficiently inhibit BChE to impart their beneficial effects.

**Table 2:** Binding energies (kcal/mol), RMSD values, and Inhibition constant (Ki) of selected ligands with PDB: 1P0I (Butyrylcholinesterase)

| Ligands | Docking Score (kcal/mol) | Reference RMSD | Estimated Inhibition Constant (Ki) |
| --- | --- | --- | --- |
| $\beta$ Carotene | -9.66 | 0.88 | 83.70 nM |
| Dihydrotanshinone I | -9.08 | 0.00 | 222.74 nM |
| Donepezil | -10.20 | 0.47 | 33.18 nM |
| Glabridin | -9.20 | 0.35 | 180.75 nM |
| Liriodenine | -8.26 | 0.27 | 884.09 nM |
| Morin | -8.57 | 0.51 | 523.04 nM |
| N-Formylanonaine | -2.88 | 0.82 | 7.79 mM |
| Quercetin | -8.24 | 0.29 | 916.32 nM |
| Quercitrin | -8.89 | 0.80 | 302.23 nM |
| Rutin | -11.36 | 0.94 | 4.73 nM |
| Vitisinol C | -10.59 | 0.41 | 17.17 nM |

The docking interaction results of PDB: 1FKN (β-Secretase) are represented in Table 3. β-secretase enzyme majorly involves the cleavage of amyloid precursor protein which ultimately forms the β amyloid plaques.^14^ β-amyloid plaque deposition is the key factor that leads to AD progression.^15^ Our results demonstrated the best docking interaction for Viticinol C (−10.05 kcal/mol) and β Carotene (−9.14 kcal/mol) that was better than the donepezil (−8.97 kcal/mol). Moreover, the docking interaction of Dihydrotanshinone I and Glabridin were comparable to standard was observed to be −8.28 kcal/mol and −8.08 kcal/mol respectively. RMSD value and Ki values for the herbal molecules were observed in the range of 0.00 to 1.00 and 42.94 nM to 7.35 mM respectively, suggesting efficient ligand-protein interaction.

**Table 3:** Binding energies (kcal/mol), RMSD values, and Inhibition constant (Ki) of selected ligands with PDB: 1FKN (β-Secretase)

| Ligands | Docking Score (kcal/mol) | Reference RMSD | Estimated Inhibition Constant (Ki) |
| --- | --- | --- | --- |
| $\beta$ Carotene | -9.14 | 0.14 | 201.08 nM |
| Dihydrotanshinone I | -8.28 | 0.00 | 846.95 nM |
| Donepezil | -8.97 | 0.76 | 264.35 nM |
| Glabridin | -8.08 | 0.47 | 01.19 μM |
| Liriodenine | -7.83 | 0.00 | 01.82 μM |
| Morin | -7.05 | 0.80 | 06.77 μM |
| N-Formylanonaine | -2.91 | 0.35 | 07.35 mM |
| Quercetin | -7.03 | 0.64 | 07.01 μM |
| Quercitrin | -7.23 | 1.00 | 04.98 μM |
| Rutin | -7.64 | 1.00 | 02.52 μM |
| Vitisinol C | -10.05 | 0.74 | 42.94 nM |

Overall, the results of the docking study suggest that Viticinol C, β Carotene, rutin, dihydrotanshinone I, and glabridin are having a strong potential to interact with AChE, BChE, and β-Secretase and thereby could exploit these pathways to impart beneficial effects during AD pathogenesis.

### 3.2 Effect of herbal molecules on HgCl_2_ induced neurotoxicity

HgCl_2_ is a well-known neurotoxin that is known to neuronal damage resembling that observed during AD through necrosis, apoptosis, and inflicting damage to the neuronal cytoskeletal. ^11^ In our study, we used 25 μM HgCl_2_ to inflict neuronal damage in Neuro-2a cell lines and evaluated the neuroprotective effect of herbal molecules in terms of percent cell viability. The results are depicted in Fig. 1 and the results indicate that quercetin and rutin are the most promising molecules that are capable of rescuing neurons from neurodegeneration.

**Fig 1:**
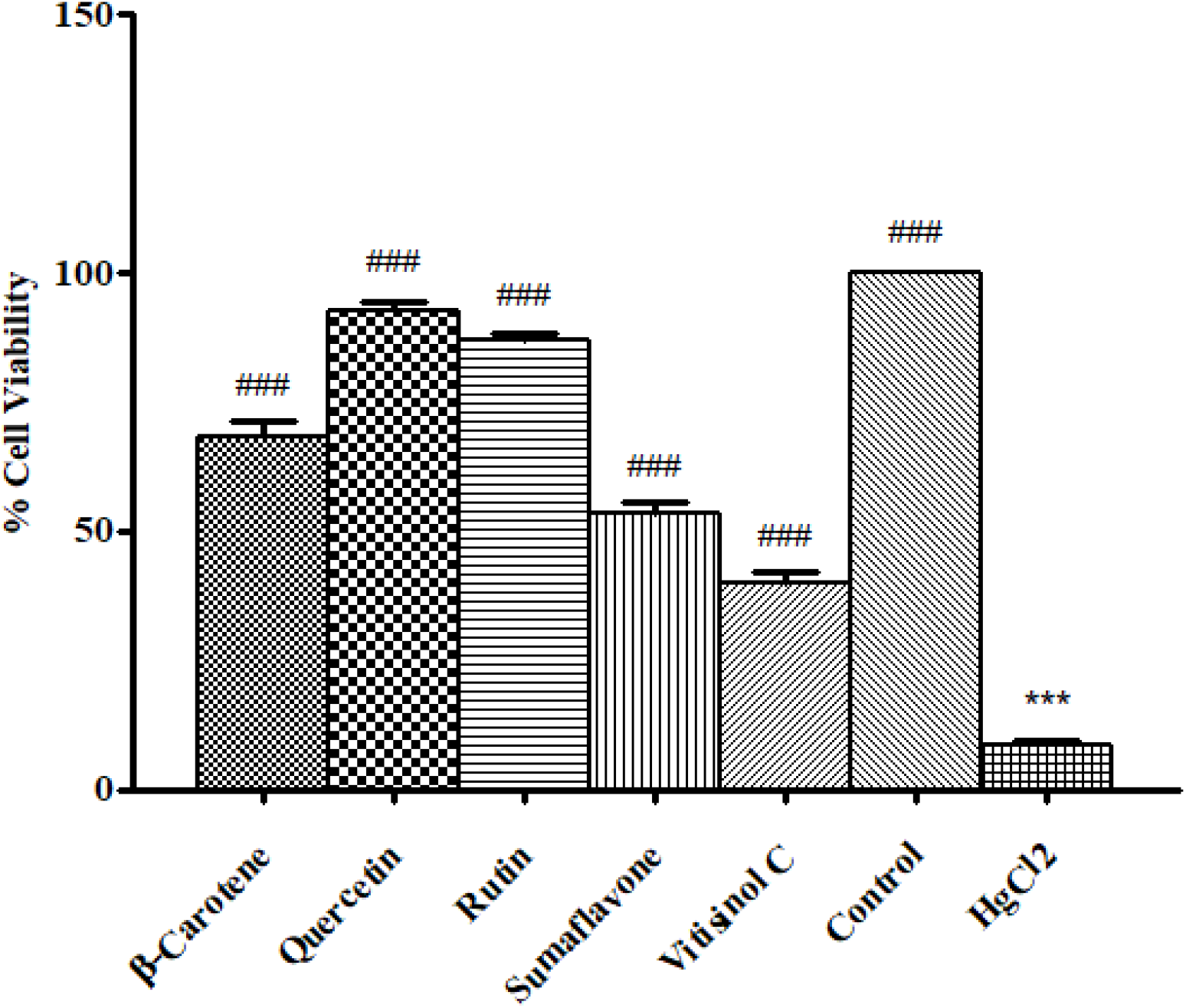
Effect of drug treatment on HgCl_2_ induced neurotoxicity calculated through MTT-assay. Results are depicted as mean ± SD (n = 3). Statistical significance was determined by one way ANOVA followed by Dunnett’s multiple comparison post hoc test at *p < 0.05, **p < 0.01 and ***p < 0.001 (HgCl_2_ v/s Control) and ^#^p < 0.05, ^##^p < 0.01 and ^###^p < 0.001 (HgCl_2_ v/s treatments).

Our results of the MTT assay demonstrated marked neuronal destruction in the cells treated with HgCl_2_ as suggested by the significantly (p < 0.001) reduced formazan formation in these cells when compared to control. The cells treated with quercetin, and rutin showed significantly (p < 0.001) higher formazan formation in the MTT assay when compared to control, which indicates a higher viable cell count in the cells treated with these molecules. Percent cell viability was observed to be 92.59 ± 2.00 for quercetin and 87.18 ± 0.92 for rutin. β-carotene, sumaflavone, and vitisinol-C also imparted significant (p < 0.001) neuroprotection against HgCl_2_ induced neurodegeneration with the percent neuroprotection of 68.42 ± 2.87, 53.56 ± 1.94, and 39.95 ± 2.09 respectively, however, the results were not as promising as those observed for quercetin and rutin. These results are in line with the previous findings where quercetin and rutin have been reported to possess good neuroprotective effects.^16^ These results suggest that quercetin and rutin can efficiently protect neurons against the neurodegenerative effect of HgCl_2_, which resembles the degenerative pattern observed during AD through cytoskeletal loss, necrosis, and apoptosis^17^, and thereby could be efficient in protecting neurodegeneration during AD.

### 3.3 Effect of herbal molecules on AChE activity

Targeting AChE is one of the most promising approaches utilized in the clinical setting for the management of AD.^18^ *In-vitro* inhibition of AChE activity is one of the most widely used methods to screen molecules with the potential to target the AChE enzyme. It is based on the detection of the formation of 5-mercapto-2-nitrobenzoic acid spectrophotometrically as a result of the reaction between DTNB and thiocholine (formed as the result of AChE mediated hydrolysis of the the substrate).^19^ The results of the effect of herbal molecules on AChE activity are demonstrated in Fig. 2(A) in terms of IC_50_ values. Our results demonstrated donepezil as the most potent molecule to inhibit AChE activity as indicated by the IC_50_ value of 39.62 nM. Donepezil is a well-known inhibitor of AChE and our results agree with the previous findings where donepezil has demonstrated strong inhibition of AChE activity in *in-vitro* settings.^20^ Our results demonstrated quercetin and rutin to be the most potent herbal molecules to inhibit AChE activity and the IC_50_ values for these molecules were observed to be 192.96 ± 17.54 μM and 335.07 ± 21.12 μM respectively. Although the IC50 values for these molecules are significantly higher than donepezil, still having IC_50_ values in this range suggests these molecules be a strong inhibitor of AChE activity. These findings are in line with previous findings where quercetin and rutin have been reported to possess the good potential to inhibit AChE activity in *in-vitro*^21, 22^, pre-clinical settings^23^, and clinical settings.^24^ β-carotene, sumaflavone, and vitisinol-C also resulted in a dose-dependent inhibition of AChE activity, and their IC_50_ values (508.60 ± 43.10, 694.18 ± 31.58, and 1009.87 ± 136.30 respectively) were observed to be higher than quercetin and rutin. These results suggest that quercetin and rutin could find an application in AD management by inhibiting AChE as inhibitors of AChE are known to slow down the progression of AD and are efficient in delaying AD onset.^25^ Further, both of these molecules are excellent antioxidants^26^ and therefore could impart additional benefit in AD pathogenesis by reducing the oxidative stress that is one of the critical pathways leading to neurodegeneration during AD.

**Fig 2:**
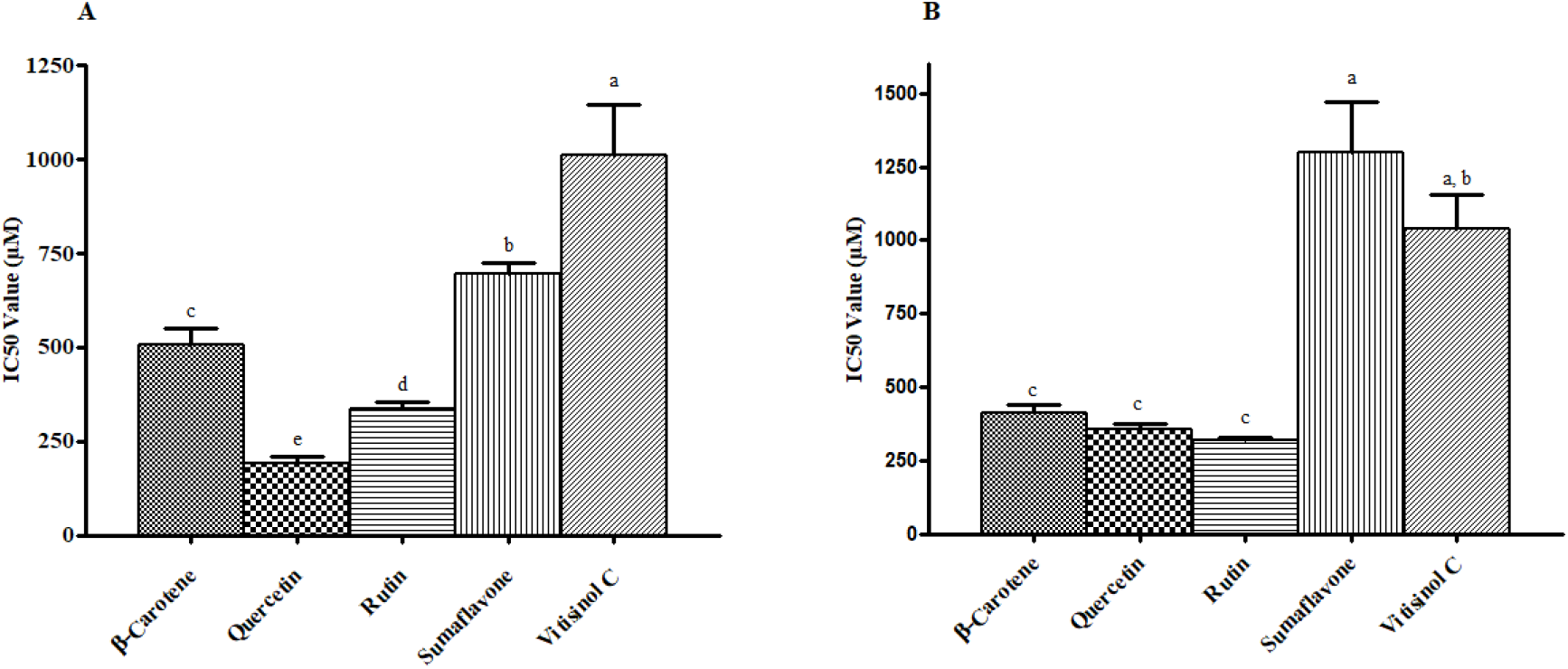
Inhibitory Concentration (IC50) value of test molecules against **(A)** Acetylcholinesterase enzyme (AChE) activity and **(B)** Butyrylcholinesterase (BChE) activity. Data are expressed as mean ± SD and the statistical significance was determined by Duncan’s multiple range test. Different letters above the error bar represent that results are significantly different at p < 0.05 by Duncan’s multiple range test.

### 3.4 Effect of herbal molecules on BChE activity

BChE is another enzyme that is known to affect the cholinergic signaling in the brain and has been demonstrated to play a critical role in the pathogenesis of AD.^27^ Like AChE assay, *in-vitro* inhibition of BChE activity is one of the most widely used methods to screen molecules with the potential to target the BChE enzyme. It is also based on the detection of the formation of 5-mercapto-2-nitrobenzoic acid spectrophotometrically as a result of a reaction between DTNB and thiocholine (formed as a result of BChE mediated hydrolysis of butyrylthiocholine iodide).^28^ The results of the effect of herbal molecules on BChE activity are demonstrated in Fig. 2(B) in terms of IC_50_ values. Our results demonstrated donepezil as the most potent molecule to inhibit BChE activity as indicated by the IC_50_ value of 142.52 ± 18.42 nM. Although donepezil is a well-known inhibitor of AChE, itis has good potential against the BChE enzyme. Our results are in agreement with the previous findings where donepezil has demonstrated strong inhibition of BChE activity in *in-vitro* settings.^22^ Our results demonstrated rutin, quercetin, and β-carotene to be the most potent herbal molecules to inhibit BChE activity in a dose-dependent manner and the IC_50_ values for these molecules were observed to be 321.32 ± 6.44 μM, 355.61 ± 20.37 μM, and 411.46 ± 27.65 μM respectively. Compared to donepezil, IC50 values for these molecules are higher, still, IC50 values in this range suggest these molecules be a strong inhibitor of BChE activity. These findings are in line with previous findings where rutin, quercetin, and β-carotene have been reported to possess the good potential to inhibit BChE activity in *in-vitro*^29,30^, pre-clinical settings^31^, and clinical settings ^31^. Sumaflavone, and vitisinol-C also resulted in dose-dependent inhibition of BChE activity, and their IC_50_ values (1299.95 ± 170.19 μM, and 1038.05 ± 119.37 μM respectively) were observed to be on a higher side when compared to other molecules suggesting their lower potential to inhibit BChE activity. These results suggest that rutin, quercetin, and β-carotene could find an application in AD management by inhibiting BChE as inhibitors of BChE are known to impart beneficial effects in the pathogenesis of AD.^32^ Further, rutin, quercetin, and β-carotene are excellent antioxidants^26^ and therefore could impart additional benefits in AD pathogenesis by reducing oxidative stress which is one of the critical pathways leading to neurodegeneration during AD.

## 4 Conclusion

The present study suggests that herbal molecules have the potential to target multiple pathways that contributes to the pathogenesis of AD and therefore could be beneficial in managing the development and progression of AD. Docking screening of the herbal molecules against AChE, BChE, and β-Secretase suggests that quercetin, rutin, sumaflavone, vitisinol-C, and β-carotene are having good affinity against these targets and the findings were further consolidated by the RMSD values and Ki values of these molecules, which were comparable to the internal standard. These results suggest that these molecules could be beneficial in managing AD pathogenesis via interfering with AChE, BChE, and β-Secretase pathways during AD. The results of the *in-vitro* neurodegeneration studies demonstrated quercetin and rutin to be the most potent molecules to rescue neurons against HgCl_2_-induced neurodegeneration. Moreover, quercetin and rutin were efficient in inhibiting AChE and BChE enzymatic activity *in-vitro*, suggesting that these molecules could modulate cholinergic pathways during AD and thereby could impart beneficial effects during AD. However, the findings demonstrate in the present study are preliminary and need to be further justified extensively through preclinical investigation to reach a decisive conclusion about their potential against AD pathogenesis.

## 5. Acknowledgement

The authors would like to acknowledge SJJT University, Jhunjhunu, and Govt. College of Pharmacy, Rohru to provide us with the facility to carry out this research work.

