## Supplemental Table 1 for "Screening of herbal molecules for the management of Alzheimer’s Disorder through molecular docking and *in-vitro* investigation"

| **Herbal molecules** | **1EVE** |
| --- | --- |
| Sumaflavone | TYR121 TRP84 TYR334 PHE331 PHE330 GLU199 |
| Rutin | PHE331 TYR334 GLY118 TYR334 PHE330 TRP84 GLU199 |
| Glabridin | TYR121 GLY118 PHE331 PHE330 TRP84 GLY118 |
| Quercitrin | TYR121 PHE331 TRP84 TYR334 PHE330 |
| Beta Carotene | PHE331 PHE330 TYR334 TRP84 GLY118 TYR121 |
| Vitisinol C | PHE330 TYR334 PHE331 ILE287 |
| N-formylanonaine | TRP84 PHE330 |
| Donepezil | GLU199 PHE331 PHE330 TRP84 TYR334 ILE287 TYR121 GLY118 |
| Dihydrotanshinone I | PHE331 PHE330 TYR334 |
| Liriodenine | TYR121 TYR334 PHE330 TRP84 |
| Quercetin | GLY118 TYR121 PHE331 |
| Morin | TYR121 TRP84 PHE330 PHE331 |

| **Herbal molecules2** | **4B0P** |
| --- | --- |
| Quercitrin | TYR332 TRP82 THR120 |
| Vitisinol C | THR120 TRP82 TYR332 |
| Rutin | TYR332 TRP82 ASP70 THR120 |
| N-formylanonaine | THR120 TYR332 TRP82 |
| Glabridin | ASP70 TYR332 TRP82 |
| Dihydrotanshinone I | TRP82 THR120 TYR332 |
| Beta Carotene | TRP82 ASP70 THR120 TYR332 |
| Morin | TYR332 TRP82 |
| Quercetin | TRP82 |
| Donepezil | TRP82 THR120 TYR332 ASP70 |
| Sumaflavone | TYR332 TRP82 THR120 ASP70 |
| Liriodenine | ASP70 TRP82 THR120 |

| **Herbal molecules3** | **1J1B** |
| --- | --- |
| Dihydrotanshinone I | ILE562 LEU688 LYS585 VAL570 |
| Quercitrin | GLN685 GLY565 GLY568 VAL570 |
| Glabridin | LYS585 GLY565 PHE567 VAL570 |
| Liriodenine | ALA583 LEU688 LYS585 VAL635 |
| Rutin | ALA583 ILE562 LEU688 VAL635 |
| Sumaflavone | THR638 VAL635 |
| Vitisinol C | ILE562 GLN685 THR638 |
| N-Formylanonaine | ALA583 LEU688 VAL635 |
| Beta Carotene | ALA583 GLN685 GLY568 LYS585 PHE567 VAL570 VAL635 |
| Quercetin | LYS585 VAL570 |
| Morin | LYS585 VAL570 |
| Donepezil | GLN685 GLY565 GLY568 ILE562 LEU688 LYS585 PHE567 THR638 VAL570 VAL635 |

| 1J1B | Docking Score  (kcal/mol) | RMSD from  reference structure | rmstol  A | Estimated Inhibition Constant, Ki |
| --- | --- | --- | --- | --- |
| β Carotene | **-9.1** | **23.78** | **0.94** | **9.63 uM** |
| Dihydrotanshinone I | **-10.1** | **19.75** | **0.45** | **1.63 uM** |
| Donepezil | **-8** | **21.55** | **1.00** | **27.59 uM** |
| Glabridin | **-9.6** | **41.62** | **0.94** | **2.52 uM** |
| Liriodenine | **-9.6** | **17.56** | **0.59** | **6.45 uM** |
| Morin | **-8.4** | **27.81** | **0.80** | **2.95 uM** |
| N-Formylanonaine | **-9.2** | **51.94** | **0.45** | **6.84 uM** |
| Quercetin | **-8.6** | **26.66** | **1.00** | **832.25 nM** |
| Quercitrin | **-9.9** | **16.39** | **0.86** | **3.82 uM** |
| Rutin | **-9.6** | **40.06** | **0.94** | **49.46 uM** |
| Sumaflavone | **-9.6** | **27.79** | **1.00** | **11.17 mM** |
| Vitisinol C | **-9.5** | **52.9** | **0.94** | **616.65 nM** |

| 4B0P | Docking Score (kcal/mol) | RMSD from reference structure A | rmstol  A | Estimated Inhibition Constant, Ki |
| --- | --- | --- | --- | --- |
| β Carotene | **-10** | **40.77** | **0.94** | **80.76 nM (nanomolar)** |
| Dihydrotanshinone I | **-10.1** | **52.87** | **0.51** | **7.29 uM (micromolar)** |
| Donepezil | **-9.7** | **41.63** | **1.00** | **30.95 uM (micromolar)** |
| Glabridin | **-10.2** | **33.44** | **0.41** | **3.43 uM (micromolar)** |
| Liriodenine | **-11.1** | **53.59** | **0.65** | **2.85 uM (micromolar)** |
| Morin | **-9.8** | **52.76** | **0.76** | **5.30 uM (micromolar)** |
| N-formylanonaine | **-10.4** | **50.33** | **0.51** | **3.00 uM (micromolar)** |
| Quercetin | **-9.8** | **52.49** | **0.82** | **6.21 uM (micromolar)** |
| Quercitrin | **-12.1** | **47.48** | **0.86** | **1.60 uM (micromolar)** |
| Rutin | **-11.2** | **44.11** | **0.76** | **67.50 uM (micromolar)** |
| Sumaflavone | **-12** | **32.42** | **1.00** | **24.81 uM (micromolar)** |
| Vitisinol C | **-11.4** | **48.94** | **0.94** | **390.25 nM (nanomolar)** |

| 1EVE | Docking Score (kcal/mol) | RMSD from reference structure A | rmstol  A | Estimated Inhibition Constant, Ki |
| --- | --- | --- | --- | --- |
| β Carotene | **-11.7** | **94.48** | **0.76** | **9.33 nM (nanomolar)** |
| Dihydrotanshinone I | **-11** | **93.55** | **0.64** | **1.75 uM (micromolar)** |
| Donepezil | **-11.2** | **93.17** | **1.00** | **1.43 uM (micromolar)** |
| Glabridin | **-10.2** | **88.47** | **0.47** | **65.49 nM (nanomolar)** |
| Liriodenine | **-10.9** | **91.75** | **0.28** | **3.26 uM (micromolar)** |
| Morin | **-10.4** | **89.95** | **0.14** | **1.88 uM (micromolar)** |
| N-formylanonaine | **-11.4** | **91.6** | **0.47** | **2.02 uM (micromolar)** |
| Quercetin | **-10.5** | **92.55** | **0.46** | **1.46 uM (micromolar)** |
| Quercitrin | **-11.8** | **74.66** | **0.76** | **4.89 uM (micromolar)** |
| Rutin | **-12.4** | **70.81** | **0.94** | **1.10 uM (micromolar)** |
| Sumaflavone | **-13.8** | **91.62** | **0.94** | **3.06 uM (micromolar)** |
| Vitisinol C | **-11.6** | **84.58** | **0.71** | **94.78 nM (nanomolar)** |

| Ligands | MW | logP | AlogP | HBA | HBD | TPSA | AMR | nRB | nAtom | RC | nRigidB | nAromRing | nHB |
| --- | --- | --- | --- | --- | --- | --- | --- | --- | --- | --- | --- | --- | --- |
| β Carotene | 536.44 | 14.734 | 8.935 | 0 | 0 | 0 | 189.29 | 10 | 96 | 2 | 31 | 0 | 0 |
| Dihydrotanshinone I | 278.09 | 2.572 | 1.39 | 3 | 0 | 43.37 | 84.57 | 0 | 35 | 0 | 4 | 24 | 3 |
| Donepezil | 379.21 | 2.633 | 0.364 | 4 | 0 | 38.77 | 115.79 | 6 | 57 | 4 | 25 | 2 | 4 |
| Glabridin | 324.14 | 2.266 | 0.981 | 4 | 2 | 58.92 | 99.71 | 1 | 44 | 0 | 4 | 26 | 6 |
| Liriodenine | 275.06 | 1.235 | -0.345 | 4 | 0 | 47.89 | 83.48 | 0 | 30 | 5 | 25 | 3 | 4 |
| Morin | 302.04 | 1.405 | -1.244 | 7 | 5 | 127.45 | 83.44 | 1 | 32 | 3 | 23 | 2 | 12 |
| N-Formylanonaine | 293.11 | 1.601 | 0.208 | 4 | 0 | 38.77 | 88.16 | 1 | 37 | 5 | 25 | 2 | 4 |
| Quercetin | 302.04 | 1.834 | -1.244 | 7 | 5 | 127.45 | 83.44 | 1 | 32 | 3 | 23 | 2 | 12 |
| Quercitrin | 448.1 | 0.802 | -2.835 | 11 | 7 | 186.37 | 114.76 | 3 | 52 | 4 | 32 | 2 | 18 |
| Rutin | 610.15 | -0.735 | -4.581 | 16 | 10 | 265.52 | 147.17 | 6 | 73 | 5 | 41 | 2 | 26 |
| Sumaflavone | 535.94 | 2.807 | -1.403 | 11 | 0 | 52.6 | 159.47 | 3 | 41 | 6 | 43 | 4 | 11 |
| Vitisinol C | 428.16 | 2.702 | 2.028 | 5 | 4 | 97.99 | 136.77 | 4 | 56 | 4 | 31 | 3 | 9 |

| Antioxidant assay | | | |
| --- | --- | --- | --- |
| IC_50_ Value  (µM) | **DPPH radical**  **scavenging assay** | **ABTS radical scavenging assay** | **NO radical**  **scavenging assay** |
| β Carotene | 0.86 ± 0.12 | 3.93 ± 0.06 | 3.94 ± 0.08 |
| Dihydrotanshinone I | 1.65 ± 0.15 | 4.73 ± 0.11 | 4.34 ± 0.05 |
| Donepezil | 20.20 ± 3.52 | 20.29 ± 0.25 | 15.69 ± 0.47 |
| Glabridin | 8.15 ± 1.25 | 3.75 ± 0.05 | 3.80 ± 0.07 |
| Liriodenine | 4.57 ± 0.26 | 4.23 ± 0.06 | 4.92 ± 0.23 |
| Morin | 0.59 ± 0.01 | 3.94 ± 0.07 | 4.07 ± 0.27 |
| N-Formylanonaine | 4.36 ± 0.30 | 3.28 ± 0.05 | 3.29 ± 0.03 |
| Quercetin | 0.38 ± 0.01 | 1.38 ± 0.04 | 1.35 ± 0.08 |
| Quercitrin | 0.45 ± 0.01 | 2.36 ± 0.03 | 2.67 ± 0.13 |
| Rutin | 0.45 ± 0.02 | 2.45 ± 0.02 | 2.56 ± 0.09 |
| Sumaflavone | 0.39 ± 0.01 | 2.22 ± 0.13 | 2.72 ± 0.03 |
| Vitisinol C | 0.51 ± 0.02 | 3.73 ± 0.03 | 3.72 ± 0.01 |

| Anti-inflammatory Assay | | |
| --- | --- | --- |
| IC_50_ Value (µM) | **Heat Induced Hemolysis** | **Albumin denaturation** |
| β Carotene | 7.61 ± 0.16 | 7.59 ± 0.29 |
| Dihydrotanshinone I | 8.03 ± 0.43 | 7.43 ± 0.27 |
| Donepezil | 115.96 ± 8.52 | 30.24 ± 1.65 |
| Glabridin | 7.99 ± 0.32 | 7.60 ± 0.26 |
| Liriodenine | 7.26 ± 0.26 | 6.44 ± 0.20 |
| Morin | 7.08 ± 0.20 | 6.61 ± 0.01 |
| N-Formylanonaine | 8.61 ± 0.15 | 7.38 ± 0.17 |
| Quercetin | 2.71 ± 0.12 | 3.54 ± 0.20 |
| Quercitrin | 6.38 ± 0.39 | 5.53 ± 0.12 |
| Rutin | 1.89 ± 0.16 | 2.69 ± 0.12 |
| Sumaflavone | 6.30 ± 0.03 | 6.57 ± 0.24 |
| Vitisinol C | 6.06 ± 0.04 | 6.82 ± 0.41 |
